# Developmental chromatin restriction of pro-growth gene networks acts as an epigenetic barrier to axon regeneration in cortical neurons

**DOI:** 10.1101/259408

**Authors:** Ishwariya Venkatesh, Vatsal Mehra, Zimei Wang, Ben Califf, Murray G. Blackmore

## Abstract

Axon regeneration in the central nervous system is prevented in part by a developmental decline in the intrinsic regenerative ability of maturing neurons. This loss of axon growth ability likely reflects widespread changes in gene expression, but the mechanisms that drive this shift remain unclear. Chromatin accessibility has emerged as a key regulatory mechanism in other cellular contexts, raising the possibility that chromatin structure may contribute to the age-dependent loss of regenerative potential. Here we establish an integrated bioinformatic pipeline that combines analysis of developmentally dynamic gene networks with transcription factor regulation and genome-wide maps of chromatin accessibility. When applied to the developing cortex, this pipeline detected overall closure of chromatin in sub-networks of genes associated with axon growth. We next analyzed mature CNS neurons that were supplied with various pro-regenerative transcription factors. Unlike prior results with SOX11 and KLF7, here we found that neither JUN nor an activated form of STAT3 promoted substantial corticospinal tract regeneration. Correspondingly, chromatin accessibility in JUN or STAT3 target genes was substantially lower than in predicted targets of SOX11 and KLF7. Finally, we used the pipeline to predict pioneer factors that could potentially relieve chromatin constraints at growth-associated loci. Overall this integrated analysis substantiates the hypothesis that dynamic chromatin accessibility contributes to the developmental decline in axon growth ability and influences the efficacy of pro-regenerative interventions in the adult, while also pointing toward selected pioneer factors as high-priority candidates for future combinatorial experiments.

## INTRODUCTION

Recovery from injury to the central nervous system (CNS) is hampered by the inability of most axons to regenerate (Blackmore, 2012; Geoffroy et al., 2016; He and Jin, 2016). One reason for this failure is that many mammalian CNS neurons undergo a developmental decline in their intrinsic capacity for axon growth, reflecting both a sharp decrease in the expression of pro-growth genes across age and a corresponding increase in the expression of growth inhibitory genes (Moore et al., 2009; Liu et al., 2011; Sun et al., 2011; Du et al., 2015; Simpson et al., 2015; He and Jin, 2016; O’Donovan, 2016). To guide ongoing efforts to reprogram adult neurons back to a regeneration-competent state, it is important to clarify the intrinsic cellular mechanisms that underlie these developmental shifts in gene expression.

Despite much progress, significant gaps in knowledge remain. First, although transcriptional profiling datasets have identified a number of growth-relevant genes that change with age, we lack understanding of how genes act collectively as a system to influence axon growth ability. This system-level perspective is needed to clarify the underlying biology, and to help identify underlying regulatory mechanisms that can be manipulated for therapeutic gain. It is clear that transcription factors play an important role in orchestrating large scale changes in gene expression, but our knowledge of the pertinent factors and how to best harness them for regenerative gain is still developing (Blackmore et al., 2012; Belin et al., 2015; Wang et al., 2015a; Norsworthy et al., 2017). Finally, and perhaps most notably, the potential influence of chromatin structure on gene expression and axon growth ability in developing neurons remains unclear. In general, gene expression is controlled by at least two regulatory levels: the *trans* level, meaning the availability of relevant TFs, and the *cis* level, meaning the local chromatin environment that can act to either facilitate or prevent TF interaction. Chromatin accessibility is now established as a critical mediator of gene regulation in various processes including cell differentiation, stem cell reprogramming, activity-dependent gene expression, nervous system development etc. (Ambigapathy et al., 2015; Frank et al., 2015; Gray et al., 2017; Su et al., 2017). Interestingly, regeneration-competent DRG neurons respond to injury by increasing chromatin accessibility around pro-regenerative gene-loci, hinting that chromatin accessibility may influence axon growth ability (Puttagunta et al., 2014). It remains unclear, however, whether chromatin accessibility is similarly dynamic in developing CNS neurons and how this may impact axon growth.

To better understand these core mechanisms – gene network activity, transcription factor regulation, and chromatin accessibility - we have developed an integrated bioinformatics workflow that considers all three and probes their interactions. Drawing on a combination of publicly available datasets and our own gene profiling data, we applied this integrated workflow to cortical neurons across postnatal development to achieve new insights into the developmental decline in regenerative ability. This approach identified subnetworks of genes that change in expression during postnatal development and detected strongly correlated changes in chromatin accessibility. Moreover, when combined with *in vivo* results regarding the varying ability of pro-regenerative transcription factors to effectively trigger axon growth, this workflow detected a strong correlation between the evoked growth and the degree of chromatin accessibility. Finally, we applied this toolset to the problem of identifying potential pioneer factors that may help relax chromatin at needed loci, thus pointing the way toward rational combinations of transcription factors for axon growth. These findings suggest an important role for chromatin accessibility in axon growth and illustrate the utility of a comprehensive bioinformatic perspective to understand and ultimately reverse the developmental decline in CNS regenerative ability.

## METHODS

### Cloning and Viral Production

Mouse *KLF6* (accession BC020042) was purchased from Dharmacon and *caSTAT3* was a kind gift from Dr. Jessica Lerch (Lerch et al., 2014a) and *JUN* plasmid is described in (Lerch et al., 2014b). Genes were cloned into an AAV-compatible plasmid with CMV reporter (pAAV-MCS, Stratagene) using standard PCR amplification as in (Blackmore et al., 2012). AAV production of AAV8-KLF6, AAV-JUN, AAV-STAT3 and the previously described AAV8-EBFP and AAV8-EGFP (Blackmore et al., 2012), was performed at the Miami Project Viral Vector Core.

### Cortical cell culture and RNA-seq

All animal procedures were approved by the Marquette University Institutional Animal Care and Use Committee. Cortical neurons were prepared from early postnatal (P3) Sprague Dawley rat pups (Harlan), and procedures for dissociation, transfection, immunohistochemistry, imaging, and neurite outgrowth analysis were performed as in (Simpson et al., 2015). Briefly, motor cortices were dissected, and neurons dissociated. Cells were transfected then plated at high density (300,000 cells/well) and maintained in culture for 2 days (P5) or 2 weeks (P14) at 37 °C, 5% CO2 in Enriched Neurobasal (ENB) media (10888-022-Thermofisher, Waltham,MA). All plates were pre-coated with poly-D-lysine hydrobromide (P7886-Sigma Aldrich, St.Louis, MO) followed by laminin (L2020-Sigma Aldrich, St.Louis,MO). AAV8-EBFP (titer- 5*10^6^ total units) was added to each well at the time of plating. To enrich for neurons, 100 nM 5-Fluoro-2’-deoxyuridine (FuDR) (F0503- Sigma-Aldrich, St.Louis,MO) was added 1 day post-transduction. At 3DIV, total RNA was extracted from neurons in culture using TRIzol reagent (10296028-Thermofisher, Waltham,MA) according to manufacturer’s instructions followed by DNAase-I treatment (EN052-Thermofisher, Waltham,MA). Total RNA quantification (Q32852- RNA HS assay, Qubit Fluorometer- ThermoFisher, Waltham,MA) and quality assessment (50671512-RNA nano chips, Bioanalyzer, Agilent, Santa Clara, CA) were performed according to guidelines recommended by ENCODE consortia (goo.gl/euW5t4). All RNA samples used for library construction had RIN scores >= 9. 100 ng of total RNA from three separate cultures were used to construct replicate libraries for each timepoint. The polyadenylated fraction of RNA was enriched by bead-based depletion of rRNA using TruSeq Stranded Total RNA Library Prep Kit with Ribo-Zero according to manufacturer’s instructions (RS-122-2201, Illumina technologies, San Diego,CA), and sequenced on an Illumina HiSeq 2000 (University of Miami Genomics Core) platform to an average of 40 million paired-end reads. Preparation of cells for transduction and isolation of RNA were performed consecutively and by the same individuals to alleviate batch effects in sample preparation and sequencing.

### Bioinformatics

#### RNA-Seq data analysis

Read quality post sequencing was confirmed using FASTQC package (per base sequence quality, per tile sequence quality, per sequence quality scores, K-mer content) (Brown et al., 2017). Trimmed reads were mapped to the rat reference genome [UCSC, Rat genome assembly: Rnor_6.0] using HISAT2 aligner software (unspliced mode along with–qc filter argument to remove low quality reads prior to transcript assembly) (Pertea et al., 2016). An average of 80–85% reads were present in map quality interval of >=30, and between 8–10% reads were excluded due to poor map quality (<10). Transcript assembly was performed using Stringtie and assessed through visualization on UCSC Genome Browser. Expression level estimation was reported as fragments per kilobase of transcript sequence per million mapped fragments (FPKM).

Distribution of reads across gene body was estimated using PICARD tools (http://broadinstitute.github.io/picard/). Differential gene expression analysis was performed using Ballgown software (Pertea et al., 2016).Isoforms were considered significant if they met the statistical requirement of having a corrected p-value of <0.05, FDR <0.05.

#### Gene network analyses

Network analysis on significant differentially expressed genes across age was performed on the cytoscape platform v3.5.1 (Lotia et al., 2013), integrating the following plug-ins – BiNGO, CyTransfinder, iRegulon, ClueGO, CluePedia and GeneMania (Maere et al., 2005; Bindea et al., 2009, 2013; Montojo et al., 2010; Lotia et al., 2013; Janky et al., 2014a; Politano et al., 2016), using default parameters in a custom script.

#### ATAC-Seq data analysis and assessment of chromatin accessibility

Publicly available ATAC-seq datasets generated from mouse cortex – PO (ENCODE: ENCSR310MLB) and Adult (GEO: GSE63137)(Mo et al., 2015) were used to assess changes in chromatin accessibility across age. Official ENCODE consortia ATAC-seq bioinformatics data analysis pipeline (https://goo.gl/932sxn) was used to identify significant peaks (signal over p-value) followed by genomic annotation [ChIPSeekeR package (Yu et al., 2015)] to classify distribution of reads across genomic loci. BEDTools (v2.27.0) was used to identify unique and shared promoters across age (Quinlan and Hall, 2010). Core promoter (2000 bp upstream/300 bp downstream) specific peaks were used to assess % accessibility dynamics across age. % Reads mapped onto core promoter regions of developmentally regulated genes relative to constitutively active gene promoter regions were quantified and assigned to be either open/completely accessible or closed.

#### TF binding site analysis

JUN and STAT3 target gene networks were constructed using iRegulon (Janky et al., 2014b) using default parameters. Transcription factor binding site/motif analysis on JUN and STAT3 target gene promoters was performed using opossum v3.0 software (Kwon et al., 2012a). Search parameters used were – JASPAR CORE profiles that scan all vertebrate profiles with a minimum specificity of 8 bits, conservation cut-off of 0.40, matrix score threshold of 90%, upstream/downstream of tss – 2000/300 bps, results sorted by Z-score >=10. Pioneer index (chromatin opening index) of TFs was obtained from two independent published algorithms (Sherwood et al., 2014; Lamparter et al., 2017).

#### Genomic visualization

ATAC-seq reads were visualized on the UCSC genome browser [Genome Reference Consortium Mouse Build 38 (GCA_000001635.2)/mm10].

### Viral delivery to cortical neurons and spinal injuries

All animal procedures were approved by the Marquette University Institutional Animal Care and Use Committee and complied with the National Institutes of Health Guide for the Care and Use of Laboratory Animals. Cortical neurons were transduced using intracerebral microinjection as described in (Blackmore et al., 2012; Wang et al., 2015b, 2018). Mice were anesthetized with ketamine/xylazine (100/10 mg/kg, IP), mounted in a stereotactic frame, and skull exposed and scraped away with a scalpel blade. 0.5 μl of virus particles were delivered at two sites located 0 and 0.5mm anterior, −1.3mm lateral from Bregma and at a depth of 0.55mm, at a rate of 0.05 ul/min using a pulled glass micropipette connected to a 10µl Hamilton syringe driven by a programmable pump (Stoelting QSI). After each injection, the needle was left in place for 1 minute to minimize viral solution flow upwards. Cervical dorsal hemisections were performed as in (Blackmore et al., 2012; Wang et al., 2015b). Briefly, adult female C57/Bl6 mice (>8wks age, 20-22g) were anesthetized by Ketamine/Xylazine and the cervical spinal column exposed by incision of the skin and blunt dissection of muscles. Mice were mounted in a custom spine stabilizer. Using a Vibraknife device (Zhang et al., 2004), in which a rapidly vibrating blade is controlled via a micromanipulator, a transection was made between the 4^th^ and 5^th^ cervical vertebrae, extending from the midline beyond the right lateral edge of the spinal cord, to a depth of 0.85mm.

### Immunohistochemistry

Adult animals were perfused with 4% paraformaldehyde (PFA) in 1X-PBS (15710-Electron Microscopy Sciences, Hatfield, PA), brains and spinal cords removed, and post-fixed overnight in 4% PFA. Spinal cords were embedded in 12% gelatin in 1X-PBS (G2500-Sigma Aldrich, St.Louis,MO) and cut via Vibratome to yield 80µm sagittal sections. Sections were incubated overnight with GFAP Ab (Z0334-DAKO, California, 1:500 RRID:AB_10013482), rinsed and then incubated for two hours with appropriate Alexafluor-conjugated secondary antibodies (Thermofisher, Waltham, MA, 1:500.) Fluorescent Images were acquired by an Olympus IX81/ Zeiss 880LSM microscopes.

### Quantification of axon growth

Axon growth was quantified from four 100µm sagittal sections of the spinal cord of each animal, starting at the central canal and extending into the right (injured) spinal tissue. The fiber index was calculated by dividing the spinal cord count by the total number of EGFP+ axons quantified in transverse sections of medullary pyramid, as in (Blackmore et al., 2012; Wang et al., 2015b, 2018). The number of EGFP+ profiles that intersected virtual lines at set distances from the injury site or midline of the spinal cord, normalized to total EGFP+ CST axons in the medullary pyramid, was quantified by a blinded observer on an Olympus IX81 microscope (Liu et al., 2010; Blackmore et al., 2012; Geoffroy et al., 2015). Exclusion criteria for spinal injuries were 1) lesion depth less than 800μm 2) axons with straight white matter trajectory distal to the lesion, and 3) EGFP+ axons in thoracic cord (too far for regenerative growth).

## RESULTS

### Network analysis reveals developmental regulation of growth-relevant gene modules

The intrinsic regenerative capacity in CNS neurons decreases during postnatal development, due to both down-regulation of pro-growth genes and upregulation of growth inhibitors across age (Moore et al., 2009). This decline in growth capacity can be modeled in cultures of primary cortical neurons, which display progressive slowing of axon extension with increasing postnatal age (Blackmore et al., 2010; Simpson et al., 2015; Venkatesh et al., 2016; Wang et al., 2018). To determine the underlying changes in gene expression we performed RNA-seq analysis, comparing early postnatal neurons maintained for either three or 14 days in culture. RNA (RNA Quality Score >9.0) was collected from 2–3 experimental replicates derived from three separate litters, and deep sequencing was performed on an Illumina platform to an average depth of 40 million reads per sample. RNA-seq experimental design, sequencing and library quality control were performed in accordance with ENCODE guidelines (RNA integrity, library quality, sequence read quality and distribution) (see methods and **Sup.Fig.1**). The Tuxedo suite of tools was used for RNA-Seq data analysis as detailed in **(Sup.Fig.1)**. Briefly, high-quality reads were aligned to the rat genome (Rnor_6.0), followed by transcript assembly using STRINGTIE and differential gene expression analysis using Ballgown software. This analysis revealed 634 genes that were differentially expressed across time (p-value<0.05, FDR<0.05), 53% of which were upregulated and 47% down-regulated **(Sup.Fig.1C)**. The complete set of differentially expressed genes (DEGs) are summarized in (**Sup.Table 1**).

One common approach to interpreting expression data is to apply gene ontologies and then test for enrichment in functional terms. Importantly, it is increasingly clear that this ontological enrichment is less informative when applied to entire gene lists, as compared to the alternative approach of preceding it with network analysis (Yon Rhee et al., 2008). In network analysis, genes are first clustered on the basis of known interactions that include direct physical interactions, shared transcriptional regulation, common regulatory pathways, and more. Once these clusters are defined, enrichment for functional terms within each cluster is tested separately. The key advantage to this approach is that it can detect functional enrichment in subnetworks that are otherwise diluted and undetected in analysis of the overall set. Using this network approach, we found that genes downregulated across age broke into 5 distinct subnetworks. Interestingly, these sub-networks were significantly enriched (p<0.05; Right-sided hypergeometric test with Bonferroni correction) for terms relevant to axon growth including neuron projection development, cytoskeleton organization, CNS development, axon guidance, cell-cell adhesion, and regulation of cell growth **(Fig. 1A)**. Genes upregulated across age broke into three sub-networks, one of which was highly enriched for functions associated with low growth capacity, including negative regulation of cell growth/proliferation/migration **(Fig. 1B)**. Notably, the largest upregulated sub-network was enriched for various aspects of synaptic transmission and formation, possibly related to the emerging concept of a functional trade-off between axon growth and synapse formation (Tedeschi et al., 2016; Tedeschi and Bradke, 2017). Overall these experiments identify genes that change in expression as neurons mature, and identify functional gene networks, both positive and negative, that may influence the developmental decline in axon growth ability.

**Fig.1.**
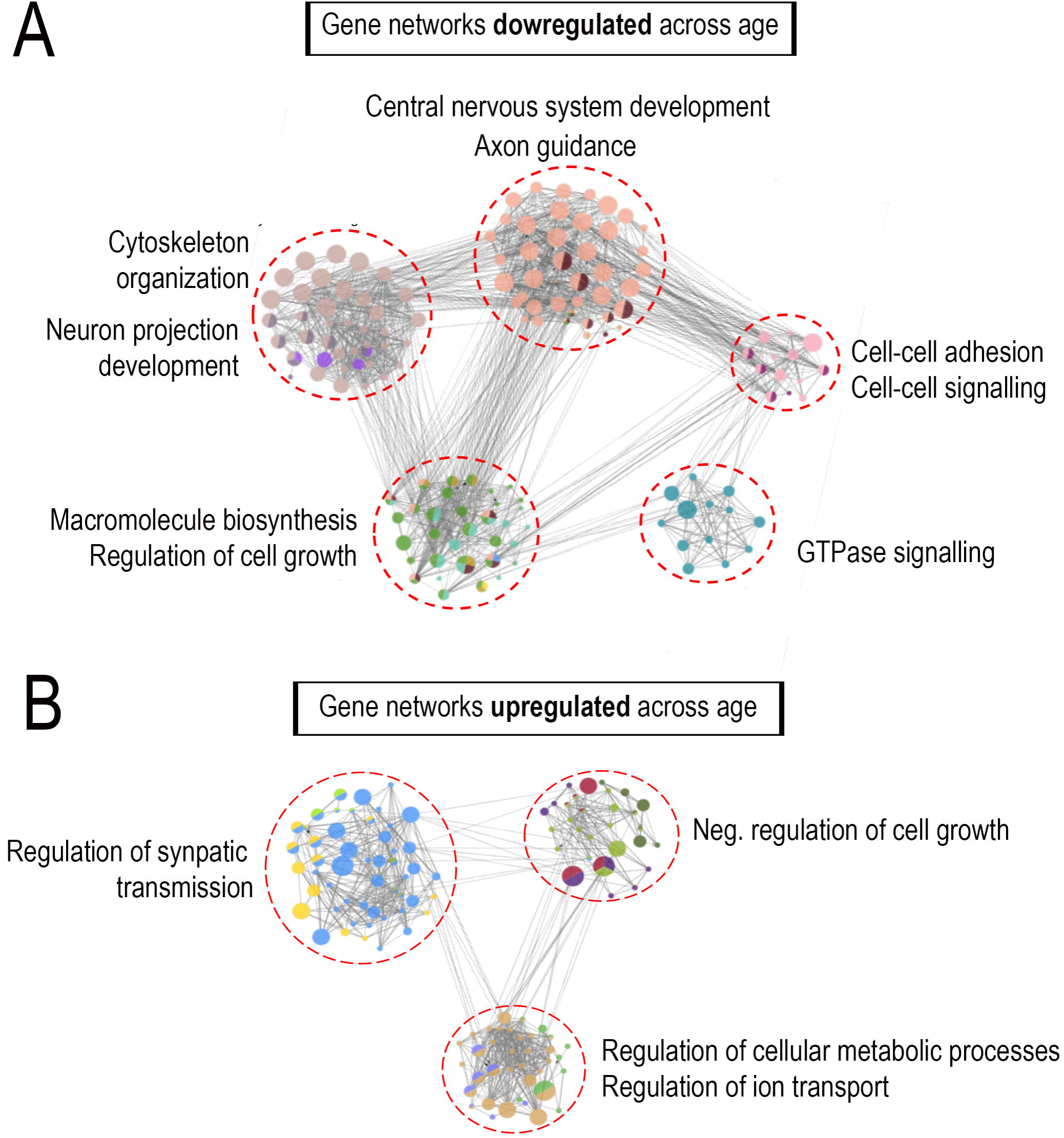
Combined RNA-Seq-network analysis reveals developmental regulation of growth-relevant gene modules. P3 cortical neurons were virally transduced to overexpress EBFP control and cultured on laminin substrates for either 2 days or two weeks before RNA extraction and deep sequencing **(A,B)** Regulatory network analysis of differentially expressed genes across age revealed sub-networks of target genes enriched for distinct functional categories highly relevant to axon growth. Nodes correspond to target genes and edges to multiple modes of interaction (physical, shared upstream regulators, shared signaling pathways and inter-regulation). Node color represents their corresponding functions as denoted in the legends and node size is based on significance for enrichment in functional category. Only significantly enriched GO categories were included in the network analysis (p<0.05; Right-sided hypergeometric test with Bonferroni correction).

### Dynamic chromatin accessibility of developmentally regulated genes

What mechanisms drive these changes in gene expression? Although the activity of transcription factors is one important and well-studied component, a second regulatory layer likely exists at the level of chromatin structure. A key measure of chromatin structure is accessibility, which describes the extent to which chromatin at a particular locus exists in an open conformation that allows interaction with regulatory proteins. Chromatin accessibility has been shown to impact numerous biological processes including nervous system development, learning and memory, cellular differentiation, stem cell biology, neurodegenerative disorders among others (Gaspar-Maia et al., 2009; Thurman et al., 2012; Ambigapathy et al., 2015; Frank et al., 2015; Gray et al., 2017; Su et al., 2017). Importantly, alterations in chromatin structure have been implicated in many aspects of human health (Gaspar-Maia et al., 2009; Hargreaves and Crabtree, 2011; Schwartzentruber et al., 2012; Fontebasso et al., 2013) motivating interest in collecting and comparing genome-wide chromatin accessibility changes that accompany various biological processes. Major collaborative research groups such as the ENCODE consortia have now generated high-quality chromatin accessibility datasets from a variety of tissues and time points (Davis et al., 2018). These derive from an ATAC-seq approach that probes DNA accessibility with a hyperactive transposase Tn5, which inserts sequencing adaptors in accessible regions (Buenrostro et al., 2015). In this way, genome-wide sequencing identifies peaks in regions of accessible chromatin. We took advantage of publicly available ATAC-seq datasets generated from mouse cortex at either post-natal day 0 (Davis et al., 2018) or adult (Mo et al., 2015) ages to identify chromatin accessibility dynamics that accompany nervous system development.

We first compared genome-wide patterns of chromatin accessibility across age. We observed that at P0, from the total accessible regions of the genome, 30% mapped to gene bodies, 20% to promoter regions, and 50% to intergenic regions (presumably enhancers) (**Fig. 2**). In adult CNS, 17% of accessible regions mapped to gene bodies, 18% to promoter regions, and 65% to intragenic/enhancer regions. Importantly, we detected a dramatic shift in the identity of accessible promoters across age. Of all promoters that were accessible at either age, only 30% were common to both, while 70% were uniquely accessible at only one age. These data indicate a large-scale, genome wide shift in the chromatin accessibility of gene promoters during the maturation of cortical tissue.

**Fig.2.**
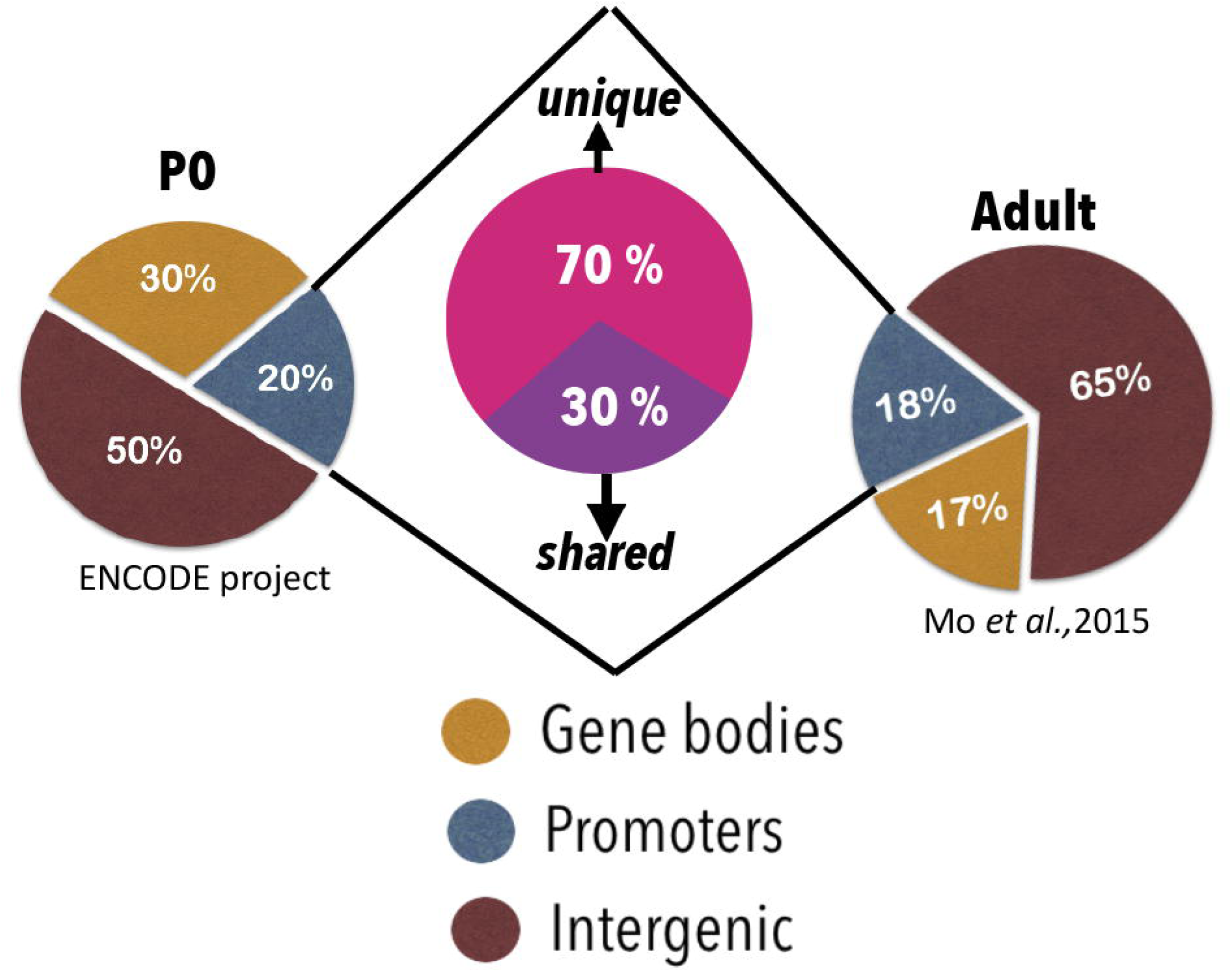
ATAC-Seq reveals genome-wide chromatin accessibility dynamics across age. Publicly available ATAC-Seq datasets generated from mouse cortex at either PO or adult ages were used to compare genome-wide changes in chromatin accessibility across age. Pie charts represent read distribution across genomic loci in P0 and adult ATAC datasets. ~18–20% gene promoters remain accessible at both time points, with 70% of the accessible promoters unique across age. ChIPseekeR package was used to annotate significant ATAC peaks (signal over p-value <0.05) to determine genomic locations corresponding to accessible peaks.

### Genes associated with axon growth become inaccessible with age

The data so far establish overall changes in both gene expression and chromatin accessibility with cortical development. As a first step in integrating these data, we examined chromatin accessibility specifically in the promoters of genes that showed differential expression across age. For each promoter region (2000 bp upstream/300 bp downstream of the transcription start site) we normalized the accessibility to within-sample housekeeping gene loci controls, and then categorized the promoter as open or closed. We looked first at the promoters of genes that were expressed at lower levels in immature neurons than in the adult (i.e. developmentally upregulated). Interestingly, only 18% of these promoters were open in younger tissue (**Fig.3A**). In contrast, 95% of these promoters were open in adult cortical tissue (**Fig.3A**). These data establish a strong correlation between promoter accessibility and the expression of developmentally upregulated genes, and hint that in immature neurons, the initially closed promoter regions may contribute to low gene expression at these loci.

**Fig.3.**
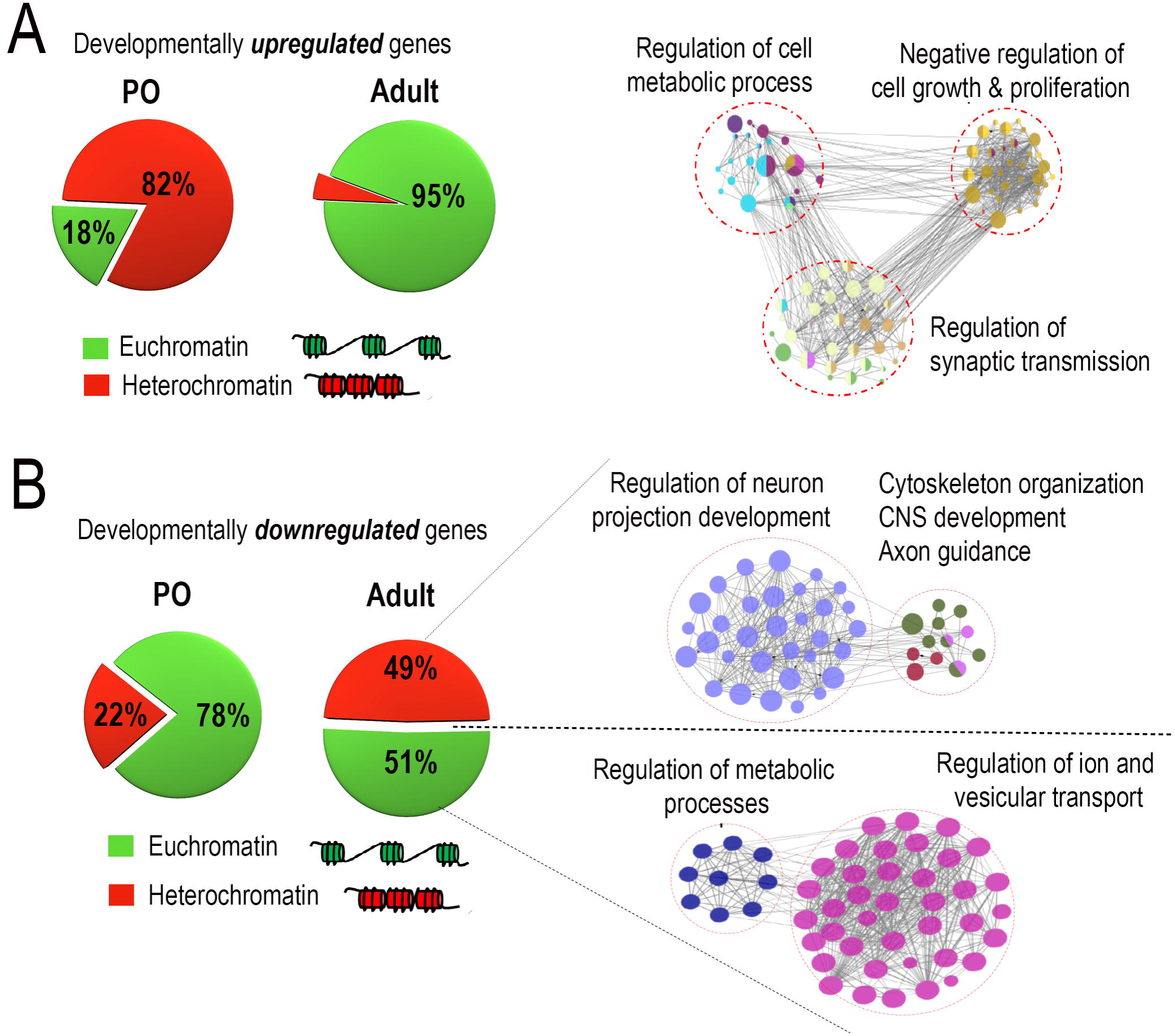
Network analysis reveals developmental regulation of growth-relevant gene modules. Chromatin accessibility around promoters of developmentally downregulated genes was assessed followed by functional classification using network analyses. **(A,B)** Pie charts represent percent chromatin accessibility around promoters of genes downregulated/Up-regulated across age (green-accessible; red – inaccessible) **(A)** Percent promoter accessibility shifts from 82% closed/ 18% open in P0 to 5% closed/ 95% open in adult neurons. Network analysis on sub-sets of genes rendered accessible across age reveal strong enrichment for GO terms negatively associated with axon growth. Network analyses were done on the cytoscape platform and only strongly enriched terms are displayed, (p-value<0.05, right-sided hypergeometric test with Bonferroni correction). **(B)** Percent promoter accessibility shifts from 22% closed/ 78% open in P0 to 49% closed/ 51% open in adult neurons. Network analysis on sub-sets of genes rendered accessible/inaccessible across age reveal preferential closure of genes enriched for GO terms highly relevant to axon growth, while genes involved in cellular homeostasis remain accessible across age

We next examined promoters of the developmentally downregulated genes. Unlike the developmentally upregulated genes, at P0 the majority (78%) of these promoters were completely accessible, while only 22% of the gene promoters were held in a closed state **(Fig. 3B)**. In adult tissue we detected a partial decline in accessibility compared to P0, such that about half of the promoters were closed and the rest remained accessible **(Fig. 3B)**. Importantly, this indicates two prominent categories of developmentally downregulated genes: those with promoters that remain accessible in the adult, and those with closed promoters. To gain insight into the functional significance of this distinction, we repeated network analysis of developmentally downregulated genes, this time dividing the genes according to their accessibility status in the adult. Interestingly, we found that the set of genes that were inaccessible in the adult were highly enriched for terms related to axon growth, including regulation of neuron projection development, cytoskeletal reorganization and axon guidance **(Fig. 3B)**. In contrast, genes that were downregulated but still remained accessible were enriched for functional categories related to cellular homeostasis, including regulation of metabolic processes and regulation of ion and vesicular transport **(Fig. 3B)**. Taken together these data indicate that cortical maturation is associated with a selective reduction of chromatin accessibility in genes associated with axon extension, suggesting a role for chromatin accessibility in restricting axon growth in the adult CNS.

### Target gene accessibility correlates with in vivo efficacy of pro-regenerative TFs

Pro-regenerative transcription factors have emerged as important tools to improve the neuron-intrinsic growth state, acting in part by activating genes that favor axon extension (Moore et al., 2009; Moore and Goldberg, 2011; Blackmore et al., 2012; Belin et al., 2015; Wang et al., 2015b; He and Jin, 2016). How might chromatin structure impact this approach? A prevalent model is that chromatin accessibility in promoter regions dictates the binding ability of transcription factors and in this way gates their influence on gene expression. If so, then simply supplying a pro-regenerative TF to an injured neuron could be insufficient to drive the expression of target genes, and thus ineffective at promoting axon growth, if chromatin compaction precludes interaction. The key prediction of this model is that pro-regenerative TFs whose target genes are accessible would be more effective in promoting axon growth than TFs whose target genes are inaccessible. To test this prediction, we selected four pro-regenerative TFs: KLF7, SOX11, JUN, and STAT3. SOX11 and KLF7 were shown previously to promote corticospinal tract regeneration when virally overexpressed *in vivo* (Blackmore et al., 2012; Wang et al., 2015b). JUN promotes axon growth when overexpressed in early postnatal cortical neurons (Lerch et al., 2014a), but its effect in adult CST neurons is unknown. For STAT3, it was recently shown that a constitutively active VP16-STAT3 mutant promotes axon regeneration in early postnatal cortical neurons (Mehta et al., 2016), but this optimized form has not been tested in the adult cortex.

Accordingly, we first tested the efficacy of forced JUN or VP16-STAT3 expression in promoting CST regeneration *in vivo*, in the same dorsal hemisection model used previously for SOX11 and KLF7 (Blackmore et al., 2012; Wang et al., 2015b, 2018). AAV8-EGFP tracer and AAV8-EBFP, AAV8-STAT3, or AAV8-JUN were delivered to the left cortex of adult female mice. One week later, CST axons in the right cervical spinal cord were transected by a dorsal hemisection injury. Animals were sacrificed eight weeks later, sagittal sections of cervical spinal cord were prepared, and GFAP immunohistochemistry was used to visualize the site of injury **(Fig. 4A-C)**. To assess axon regeneration, the number of EGFP-labeled CST axons was quantified at set distances distal to the injury), then normalized to the total number EGFP-labeled CST axons in the medullary pyramid. As expected, EBFP control animals displayed very little CST growth distal to the injury site (**Fig. 4A**). In both VP16-STAT3 and JUN-treated animals, we observed no significant elevation in CST growth compared to EBFP controls at any of the distances examined (**Fig.4 B-D**). Collectively, of four TFs with pro-regenerative properties in at least some cellular contexts, we find that two factors stimulate growth in adult CST neurons (KLF7 and SOX11), whereas two (VP16-STAT3 and JUN) stimulate do not stimulate growth in adult CST neurons at all.

**Fig.4.**
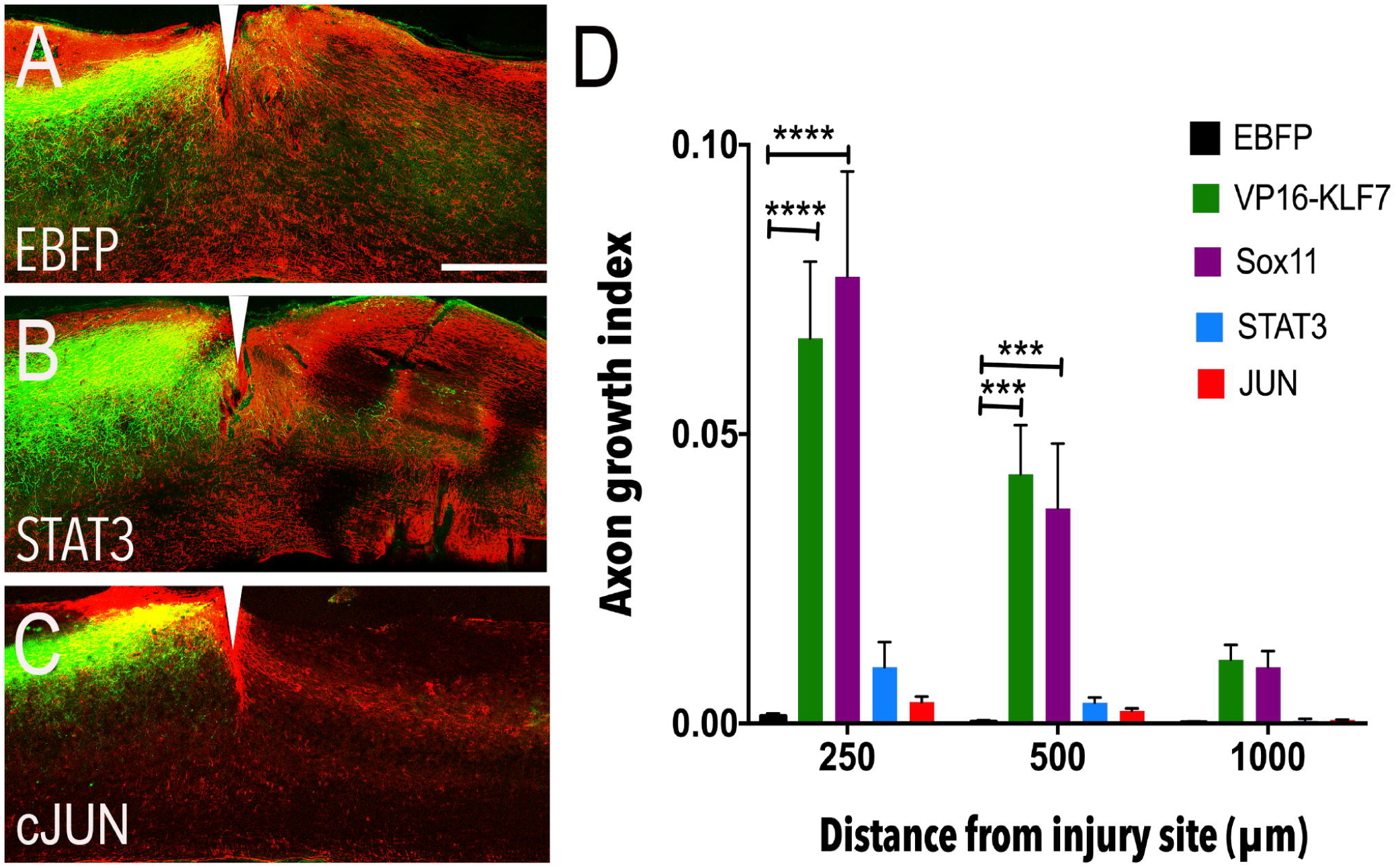
cJUN and STAT3 target gene promoters become inaccessible across age. Adult female mice received cortical injection of AAV-JUN, AAV-STAT3 or AAV-EBFP control, mixed 2:1 with AAV-EGFP tracer. One week later CST axons were severed by C4/5 dorsal hemisection. **(A-C)** Sagittal sections of spinal cord were prepared and GFAP immunohistochemistry (red) performed to define injury sites (white arrowheads). EGFP-labeled CST axons were rarely visible in tissue distal to the injury in AAV-EBFP treated animals (A), AAV-STAT3 treated animals (B) or AAV-cJUN treated animals (C) **(D)** Quantification of the number of CST axons that intersect virtual lines below the injury, normalized to total EGFP+ axons in the medullary pyramid, showed no significant elevation of axon growth with any treatment. VP16-KLF7 and Sox11 treatment values from previous publication (Blackmore et al., 2012; Wang et al., 2015a) included for reference (****p-value<0.0001; ***p-value<0.001;N=7–9 animals in each group, RM 2-way ANOVA with Sidak’s multiple comparisons test.

We next assembled lists of potential target genes for each factor, drawing on a combination of published data (Norsworthy et al., 2017), our own RNA-seq datasets, and publicly available ENCODE data (Mo et al., 2015; Davis et al., 2018). For KLF7, we used RNA-seq analysis of cortical neurons with forced overexpression of KLF6, a closely related KLF family member. KLF6 has been shown to act redundantly with KLF7 to promote regeneration, phenocopies the pro-growth effects of KLF7 in our hands, and has a nearly identical DNA binding domain (Veldman et al., 2007; Moore et al., 2009; Wang et al., 2018).We have found KLF6 protein expression to be more stable than KLF7, and for that reason have focused on this related family member in more recent work (Wang et al., 2018).Using RNA-seq analysis of cortical neurons transduced with KLF6 we assembled a list of 345 KLF6-responsive genes (Wang et al., 2018). For SOX11 we utilized a recently published dataset of neurons overexpressing SOX11 (Norsworthy et al., 2017). Finally, for STAT3 and JUN we leveraged published ENCODE ChIP-Seq data to identify loci directly bound by these TFs (see methods) (Janky et al., 2014b; Davis et al., 2018).

Next, returning to the publicly available ATAC-seq datasets, we examined the promoter accessibility of the genes in each of these four sets at both early postnatal and adult time points. At P0, all four gene sets were highly accessible: 95% of the KLF6 set, 91% of SOX11 set, 95% of JUN set, and 90% of the STAT3 set had promoters in an open conformation (**Fig.5A**). In the case of KLF6 and SOX11 target genes, this open conformation was largely maintained into adulthood, such that 90% of KLF6 and 75% of SOX11 target promoters remained open (**Fig.5A**). In striking contrast, in the JUN and STAT3 sets a much smaller fraction of target gene promoters remained accessible in adult neurons (45% of JUN target gene promoters, 40% of STAT3 target gene promoters, **Fig. 5B**). We plotted the percent accessibility of gene targets against the *in vivo* regenerative growth index at 1mm below injury sites for each factor, and found a strong correlation (**Fig. 5C**, % R^2^ = 0.965). Overall this analysis provides strong circumstantial support for the hypothesis that epigenetic constraint in the form of chromatin structure that may limit the full efficacy of pro-regenerative TF treatments.

**Fig.5.**
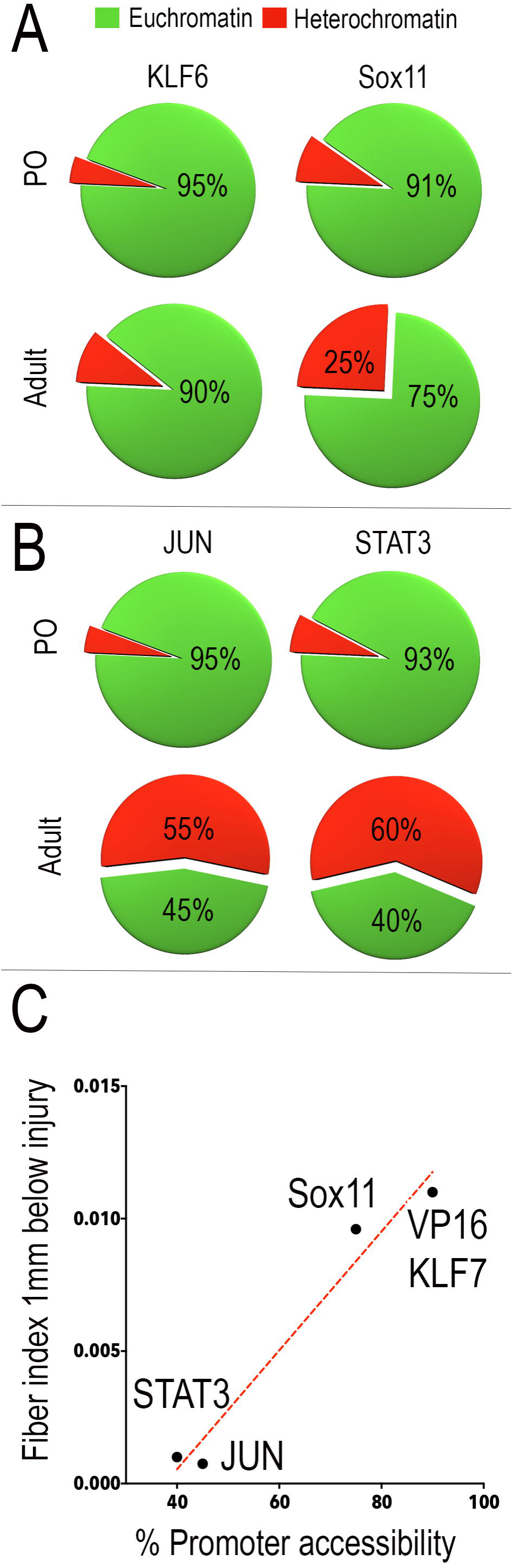
Axon growth response produced by pro-regenerative TFs correlates with accessibility of target genes. Pie charts represent percent promoter accessibility of pro-regenerative TF targets across age (green-accessible; red – inaccessible) **(A)** Promoters of KLF6 and Sox11 targets are accessible in adult neurons (KLF6 – 95% open in P0 to 90% open in adult neurons; Sox11 – 91% open in P0 to 75% open in adult neurons). **(B)** Promoters of JUN and STAT3 targets are largely inaccessible in adult neurons (JUN- 95% open in P0 to 45% open in adult neurons; STAT3- 93% open in P0 to 40% open in adult neurons). **(C)** Percent promoter accessibility of TF target genes plotted against axon growth index at 1mm from injury shows strong correlation between target gene accessibility and *in vivo* efficacy of pro-regenerative TFs. Red dotted line displays best line of fit (degrees of freedom – 2; R square = 0.965).

### Motif analysis predicts relevant combinations of pioneer factors

So far our analyses indicate that chromatin compaction may contribute to the developmental decline in pro-regenerative gene expression, and suggest that it may also limit the ability of exogenously supplied transcription factors to reactivate these genes. Beyond identifying this challenge, how might bioinformatic analyses point toward a solution? One common approach to improving accessibility involves manipulation of post-translational modifications on histones. Acetylation of histones, mediated by Histone Acetyltransferases (HATs) generally leads to a more relaxed, accessible chromatin state, whereas deacetylation mediated by Histone Deacetylases (HDACs) restricts DNA accessibility (Bannister and Kouzarides, 2011). HDAT/HAT manipulation has shown some ability to alter axon growth in PNS models of regeneration (Gaub et al., 2010; Cho and Cavalli, 2014; Puttagunta et al., 2014). Targeting histone modifications lacks specificity, however, and in CNS neurons has so far yielded minimal improvements (Venkatesh et al., 2016). As an alternative, we considered a strategy that could potentially yield more targeted chromatin remodeling and which is gaining traction in the stem cell biology field, namely the use of Pioneer TFs. Pioneer TFs are defined by virtue of their unique ability to bind closed chromatin and initiate a cascade of molecular events that leads to increased accessibility (Zaret and Carroll, 2011; Iwafuchi-Doi and Zaret, 2014; Sherwood et al., 2014; Zaret, 2018). Knowledge of TFs with pioneer activity is rapidly expanding, in part fueled by the development of computational algorithms that can predict pioneer activity in TFs based on “chromatin opening index / pioneer index of TFs” assessed from genome-wide chromatin accessibility datasets (Sherwood et al., 2014; Lamparter et al., 2017). This raises the possibility that if appropriate pioneer factors could be provided to injured neurons along with pro-regenerative TFs such as JUN or STAT3, they could potentially render promoters more accessible and thus help relieve the chromatin-based constraint on pro-regenerative efficacy.

We therefore applied our bioinformatic framework to the task of identifying potential pioneer TFs that could regulate the accessibility of JUN or STAT3 target genes. We first used motif-scanning algorithms to examine the core promoters of JUN and STAT3 target genes and identify significantly over-represented binding motifs for TFs (Z-score >=10) (Kwon et al., 2012b) **(Fig. 6A).** From the list of predicted TF motifs, we identified the subset with the highest pioneer index/chromatin index as described in (Sherwood et al., 2014; Lamparter et al., 2017) **(Fig. 6B).** This analysis predicted the TFs Arid3a, Spi1 and Nkx3-2 as TFs that could exert pioneer activity at the promoters of STAT3 target genes **(Fig. 6B)**, and CEBPA, EBF1 and FoxA1 as potential pioneers for JUN target genes. **(Fig. 6B)**. This analysis points the way toward rational combinations of TFs to combine with JUN/ STAT3 to achieve targeted chromatin remodeling and potentially enhance the efficacy of these factors in promoting regenerative ability in the cellular context of the adult cortex.

**Fig.6.**
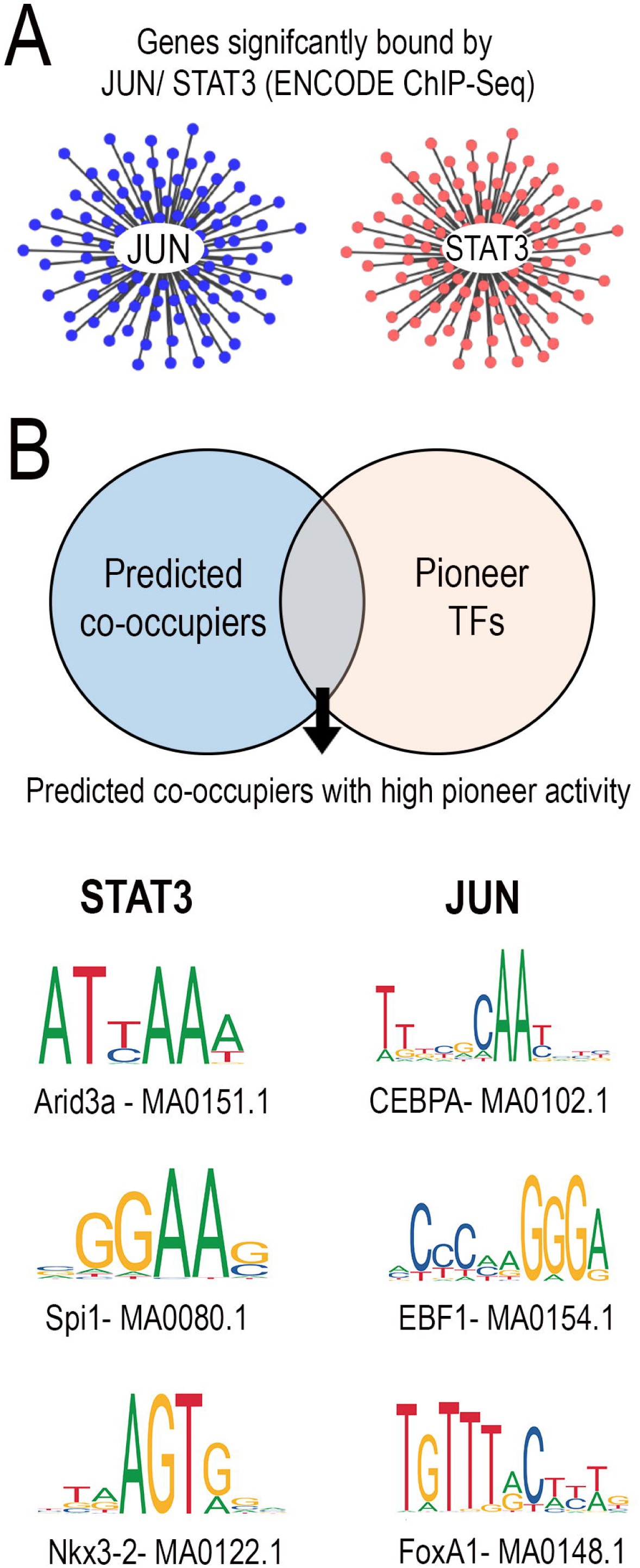
Motif analysis predicts relevant combinations of pioneer factors (A,B) JUN and STAT3 target gene networks were constructed from ENCODE ChIP-Seq data. TFs predicted to co-occupy gene promoters alongside either JUN/STAT3 were identified through motif analysis (p-value <0.0001) and compared against the list of pioneer TFs with the highest chromatin opening index. **(C)** Pioneer TFs- Arid3a, Spi1 and NKX3-2 were predicted to be relevant combinations for STAT3 and pioneer TFs- CEBPA, EBF1 and FoxA1 were predicted to be relevant pioneer TF combinations for JUN.

## DISCUSSION

To better understand the developmental loss of axon growth ability in CNS neurons we have applied an integrated bioinformatics workflow comprised of gene network analysis, TF motif analysis, and genome-wide assessment of chromatin accessibility. This analysis identified sub-networks of developmentally regulated genes that likely influence axon growth ability, which underwent selective chromatin closure in promoter regions with age. We also showed that unlike SOX11 and KLF7, VP16-STAT3 or JUN are ineffective in promoting CST regeneration in adult animals. Intriguingly, our analysis indicates that the promoters of putative target genes of STAT3 and JUN are on average less accessible than those of SOX11 or KLF7, hinting at chromatin-based constraints. Finally, the workflow predicted a set of pioneer factors that could potentially open the promoters of JUN or STAT3 target genes, thus pointing toward future combinatorial TF approaches.

### The importance of chromatin accessibility for axon growth ability

Mammalian genomes harbor millions of *cis*-regulatory elements that act as landing pads for TFs. The degree of chromatin compaction can strongly influence TF interactions, and in this way local chromatin accessibility establishes transcriptionally active versus inactive regions of the genome (Kwon and Wagner, 2007; Tsompana and Buck, 2014). Originally identified in stem cell biology, the significance of chromatin accessibility in regulation of gene expression has been recently confirmed in a variety of neurobiological processes including nervous system development (Frank et al., 2015), cortical neurogenesis (de la Torre-Ubieta et al., 2018), activity-dependent gene expression (Su et al., 2017) among others. Recently, integrated neuronal chromatin accessibility and gene expression profiles have identified vast developmental changes in chromatin accessibility that strongly correlate with changes in gene expression genome-wide (Frank et al., 2015; de la Torre-Ubieta et al., 2018).

These findings raise the possibility that changes in chromatin accessibility during neural maturation could similarly influence the expression of genes needed for effective axon growth (Trakhtenberg and Goldberg, 2012; Finelli et al., 2013; Puttagunta et al., 2014). Previously, we tested this idea by examining histone modifications near selected regeneration-associated genes, and detected age-related changes consistent with chromatin closure (Venkatesh et al., 2016). The present approach offers several key improvements that significantly strengthen the link between chromatin accessibility and regeneration-associated gene expression. First, the ATAC-Seq approach provides direct information on chromatin accessibility, as opposed to measuring histone modifications as an indirect surrogate. Second, rather than selecting specific genes for study, the current dataset enables unbiased analyses that span the entire genome. Finally, we have taken an integrated approach that combines genome-wide differential gene expression datasets and chromatin accessibility datasets, which together are far more powerful than either alone. Interestingly, our analysis indicates that not all genes that are developmentally downregulated necessarily undergo closure, but rather about half retain an open conformation in their promoter regions. One interpretation is that some genes remained poised for expression, for instance if relevant TFs become activated, while other genes are more completely silenced. Perhaps the most important finding is that the set of genes that display chromatin closure, but not the open cohort, are highly enriched for regeneration-relevant functions. This result substantiates the emerging notion that reduced chromatin accessibility presents an obstacle for the re-expression of regeneration associated genes in adult neurons (Puttagunta et al., 2014; Venkatesh et al., 2016).

An important question, however, is whether changes in chromatin accessibility are the cause or consequence of gene transcription. In other systems there is strong evidence for the primary role of accessibility, based on careful time course analyses. High-frequency datasets of accessibility and gene expression that span key stages in cellular reprogramming and/or cell differentiation have confirmed that local changes in accessibility precede and are a pre-requisite for enhanced transcriptional output (Iwafuchi-Doi and Zaret; Sherwood et al., 2014; Frank et al., 2015; Li et al., 2017; de la Torre-Ubieta et al., 2018; Zaret, 2018). Studies in regeneration-competent DRG neurons also lend support to this idea. First, transcriptional profiles across early stages of PNS regeneration have revealed a clear hierarchy in the activation of target genes following injury, starting with a wave of chromatin remodelers that precedes activation of well-established regeneration associated genes (Li et al., 2015). In further support of this, injured DRG neurons retrogradely activate specific histone acetyltransferases, a key first step, that lead to local chromatin remodeling around regeneration-associated genes culminating in transcriptional activation of these genes (Puttagunta et al., 2014). Similar time series have yet to be generated in the developing or injured CNS. Nevertheless, based on the available information from other cell types, we favor a model in which changes in accessibility during CNS maturation are not merely passive indicators of transcriptional activity, but rather contribute functionally to changes in gene expression.

### Chromatin accessibility as a predictor of TF phenotype

JUN and STAT3 have been shown previously to act as pro-regenerative factors in a variety of cellular contexts (Raivich et al., 2004; Bareyre et al., 2011; Ruff et al., 2012; Lang et al., 2013; Lerch et al., 2014a; Fagoe et al., 2015; Luo et al., 2016; Mehta et al., 2016). Both JUN and STAT3 expression correlate with regenerative ability, as they are upregulated after injury by neuronal types that successfully regenerate, including sensory neurons and zebrafish retinal ganglion cells (Broude et al., 1997; Raivich et al., 2004; Ruff et al., 2012). In addition, knockout or functional blockade of JUN or STAT3 reduces peripheral regeneration (Raivich et al., 2004; Bareyre et al., 2011) and regeneration by zebrafish RGCs (Elsaeidi et al., 2014), showing that they are required for effective axon growth. Most relevant to the current work, JUN overexpression increases the rate of axon extension in early postnatal cortical neurons (Lerch et al., 2014a; Callif et al., 2017), indicating that in this cellular context, increased JUN levels are sufficient to potentiate axon growth. Similarly, forced expression of STAT3 in injured retinal ganglion cells in rats modestly improves regeneration (Pernet et al., 2013). Forced expression of wildtype STAT3 was tested previously in rodent models of spinal injury, and produced minimal CST sprouting that extended a few hundred microns from the injury site. More recently a modified form of STAT3 was developed in which the transcriptional activation domain VP16 was added to a constitutively active form of STAT3. This VP16-STAT3 construct, but not wildtype, enhanced axon growth when overexpressed in early postnatal cortical neurons, again demonstrating the sufficiency of forced expression to boost axon growth (Mehta et al., 2016).

We therefore hypothesized that forced expression of JUN or VP16-STAT3 would promote regeneration in adult CST neurons. We found, however, that JUN had no effect on axon growth, and VP16-STAT3 only slightly increased short distance sprouting, similar the prior test of wildtype STAT3 (fiber index 0.01 at 200um in (Lang et al., 2013), 0.95 at 250um here). This is in contrast to our prior findings with overexpression of SOX11 and VP16-KLF7, both of which produced much more substantial CST regeneration. It is important to note that we previously tested SOX11 and VP16-KLF7 in an identical injury model, and indeed the experiments were performed by the same surgeon (Blackmore et al., 2012; Wang et al., 2015a). Similarly, we have used these same JUN and VP16-STAT3 constructs in postnatal cortical neurons, and have confirmed their growth-promoting properties in these cells (Callif et al., 2017). Collectively, these data enable a clear conclusion that the phenotypic effects of overexpressed JUN and VP16-STAT3 in adult CST neurons are less than those of SOX11 and VP16-KLF7, and are less than their own effects in postnatal cortical neurons.

Here we provide circumstantial evidence that this difference in growth phenotype may be explained in part by differences the chromatin accessibility of target genes. At an early postnatal timepoint, when these factors show growth-promoting properties, the predicted target genes of all four TFs are almost complete accessible. In adult cortical tissue, the great majority of SOX11 and KLF6/7 target genes remain accessible, whereas less than half of JUN and STAT3 targets are open. Thus for these four TFs, chromatin accessibility in target genes is well correlated with the growth phenotype that results from forced expression in cortical neurons. This is not to suggest that chromatin accessibility is the only relevant factor. As just two examples, JUN activity also depends on the presence of appropriate AP1 binding partners (Hai and Hartman, 2001), and STAT3’s transcriptional output is similarly affected by binding partners and by feedback inhibition from SOCS3 (Smith et al., 2009). Nevertheless, the relationship between accessibility and TF-evoked growth is striking, and has potentially important implications for the use of TFs as pro-regenerative tools. The pattern is consistent with the notion that chromatin structure may fundamentally constrain the ability of TFs to activate target genes, and in this way fundamentally constrain their efficacy as pro-regenerative tools. If so, one interesting application of our bioinformatic pipeline would be to pre-screen future candidate TFs prior to *in vivo* testing, in order to use the promoter accessibility of the target genes as an early indicator of potential efficacy.

### Pioneer factors for regeneration

If chromatin compaction limits the transcriptional reactivation of pro-growth genes, the obvious follow-up question is how this compaction can be reversed. Chromatin structure is influenced by two main classes of proteins: histone-modifying enzymes, which ultimately regulate the degree of histone/DNA association, and ATP-dependent remodeling complexes that actively restructure chromatin (Kwon and Wagner, 2007; Bannister and Kouzarides, 2011; Hargreaves and Crabtree, 2011; Tsompana and Buck, 2014). Importantly, histone modifiers and chromatin remodelers must be recruited to specific loci. One approach to relieving chromatin-based constraints to axon growth has been to manipulate histone-modifying enzymes through pharmacological or genetic means (e.g. HDAC inhibitors) (Gaub et al., 2010; Cho and Cavalli, 2014). This strategy is highly non-selective across the genome, however, and thus doesn’t account for the locus-specific nature of chromatin closure that is apparent in our data. Perhaps for this reason, this strategy has so far has yielded limited gains in axon growth (Gaub et al., 2010; Finelli et al., 2013; Puttagunta et al., 2014; Venkatesh et al., 2016). How, then, can global remodelers be directed to specific genomic loci? Work in stem cell biology has revealed that during cellular reprogramming events, a class of molecules called Pioneer TFs initially engage with compacted chromatin at specific loci, flagging these regions for subsequent chromatin relaxation and gene expression (Iwafuchi-Doi and Zaret, 2014; Sherwood et al., 2014; Morris, 2016; Guo and Morris, 2017). Thus, pioneer factor binding marks the first in a sequence of events that culminates in recruitment of chromatin remodelers and transcriptional activators. However, it is notable that in the absence of any other transcription factors, pioneer TF binding alone is insufficient to induce changes in gene expression; conversely other TFs are unable to engage silent chromatin without the assistance of pioneers (Morris, 2016; Guo and Morris, 2017). Therefore, efficient cellular reprogramming requires an optimal combination of both pioneer factors and cell-specific TFs, which act co-operatively to initiate transcriptional activation at specific genomic loci.

We speculate that a similar optimal combination of pioneer factors/pro-regenerative TFs may be needed to reverse age-related epigenetic constraints at specific pro-growth genomic loci. Interestingly, regeneration-competent DRG neurons respond to injury with re-activation of genes modules in a hierarchical manner, starting with a sharp increase in the expression of several TFs that are known to act as pioneers in other cellular contexts (Li et al., 2015). It remains speculative that these TFs exert pioneer activity in injured DRGs, and that such activity is important for subsequent regeneration, but this model is quite plausible in light of the highly conserved roles for pioneer factors in multiple biological processes (Takahashi and Yamanaka, 2006; Soufi et al., 2012, 2015; Wapinski et al., 2013; van Oevelen et al., 2015). Accordingly, we have performed motif analysis of regulatory DNA that is targeted by pro-regeneration TFs but is subject to developmental closure, and have identified high enrichment for sets of pioneer factors. This analysis motivates the hypothesis that co-expression of these factors in adult tissue could enhance the efficacy of TF overexpression, providing a foundation for future combinatorial experiments.

An important caveat to these analyses regards the diverse sources of input data. Ideally, the tools presented here would be powered by unified datasets, for example RNA-seq, ATAC-seq, and ChIP-seq, data from purified CST neurons of early postnatal and adult ages. Owing to the technical challenges of obtaining such source material, and the relative nascence of attention to these approaches in the regeneration field, these ideal datasets are unavailable or in development. This is in contrast to other fields (e.g. stem cell, cancer biology) in which creating massive genome-wide profiling data is relatively simple, and it is the development of bioinformatics platforms to synthesize and interpret the data that act as the bottleneck. The regeneration field is essentially the mirror image, in which the rapidly developing concepts and tools for genome-wide analysis can be borrowed from other fields, but are held back by a dearth of input data. Accordingly, here we drew together the best information available regarding dynamic gene expression, transcription factor targets, and chromatin accessibility in postnatal and adult cortical neurons. Nevertheless, we could not avoid some discrepancies across data sets. For example, the developmental changes in gene expression were measured during postnatal maturation, whereas ATAC-seq analyses compared postnatal to fully adult tissue. In addition, the ATAC-seq data were based on a mix of whole tissue (P0 time point) and purified neurons (adult). Finally, lists of TF target genes came from RNA-seq analysis of TF overexpression and from ChIP-Seq data from mixed tissue sources, and likely only approximate the direct targets of these factors. As improved input data become available it will be essential to continually update this workflow and re-evaluate conclusions. Nevertheless, although imperfect, this approach yielded provocative results and point toward a central role for chromatin accessibility in regulating axon growth.

**Sup. Fig.1 RNA-seq library quality control and differential gene expression analyses.** P3 cortical neurons were virally transduced to overexpress EBFP control and cultured on laminin substrates for either 2 days or two weeks before RNA extraction and deep sequencing. **(A)** RNA-Seq data analysis workflow. **(B)** High mapping quality of sequenced reads confirmed library and sequencing quality. Sequencing read distribution across the gene body for treatment groups were monitored to ensure optimal library and sequencing quality. **(C)** Differential gene expression analysis identified 634 genes that significantly differ between P5 and P14 groups (p-value<0.05, FDR<0.05). 53% of the transcripts were upregulated and 47% of the transcripts were down-regulated across age.

## Acknowledgments

This work was supported by grants from NINDS, the International Spinal Research Trust, the Bryon Riesch Paralysis Foundation and The Craig Nielsen Foundation. The authors declare no competing financial interests. The authors acknowledge ENCODE consortia and The Bing Ren lab at UCSD that generated the ATAC-seq datasets used in this study.

